# Phenotyping technology for assessing protein content in seaweed by field spectroscopy and a machine learning algorithm

**DOI:** 10.1101/2022.04.27.489785

**Authors:** Niva Tadmor Shalev, Andrea Ghermandi, Dan Tchernov, Eli Shemesh, Alvaro Israel, Anna Brook

## Abstract

Determining seaweed protein concentration and the associated phenotype is critical for food industries that require precise tools to moderate concentration fluctuations and attenuate risks. Algal protein extraction and profiling have been widely investigated, but content determination involves a costly, time-consuming, and high-energy, laboratory-based fractionation technique. The present study examines the potential of field spectroscopy technology as a precise, high-throughput, non-destructive tool for on-site detection of red seaweed protein concentration. By using information from a large dataset of 144 *Gracilaria* sp. specimens, studied in a land-based cultivation set-up, under six treatment regimes during two cultivation seasons, and an artificial neural network, machine learning algorithm and diffuse visible–near infrared reflectance spectroscopy, predicted protein concentrations in the algae were obtained. The prediction results were highly accurate (R^2^ = 0.95; RMSE = 0.84), exhibiting a high correlation with the analytically determined values. External validation of the model derived from a separate trial, exhibited even better results (R^2^ = 0.99; RMSE = 0.45). This model, trained to convert phenotypic spectral measurements and pigment intensity into accurate protein content predictions, can be adapted to include diversified algae species and usages.

**Highlight:** Non-destructive determination of protein content in the edible red seaweed *Gracilaria* sp. by in-situ, VIS-NIR spectroscopy and a machine learning algorithm.

## 1. Introduction

In the context of climate change and the degradation of natural resources, there is growing interest in developing novel protein sources for food security. Numerous authors have described the potential advantages of producing protein from marine sources, particularly algae (both micro and macro), in reducing environmental pressures derived from terrestrial agriculture while obtaining products with high-quality nutritional properties. (EU initiative, 2022; Gephart et al.,_2021).

Seaweeds are known for their high yields of potentially edible, high-quality protein in proportion to their dry weight (Bleakley and Hayes, 2017; Kazir et al., 2019). Seaweeds are also regarded as an important source of nutrients, vitamins, minerals, and trace elements, with broad commercial applications in food, feed, fertilizers, pharmaceuticals, nutraceuticals, and cosmetics (Charoensiddhi et al., 2020) and are considered to have potential environmental benefits (Lange et al., 2020). Numerous methods have been developed for the optimizing algae cultivation, enhancing functional traits, and reducing variability in the concentrations of functional traits, but all these methods are limited in that they are time-consuming, lack precision and are dependent on laboratory measurements. In addition, although profiling and extraction of algal proteins have been widely investigated, there is a marked a lack of progress in advancing the protein determination technology, which still involves a costly, time-consuming and high-energy laboratory-scale fractionation technique (Cermeño et al., 2020). A promising research direction is based on field spectroscopy, which has already been used successfully for diversified applications in precision agriculture (Lee and Ehsani, 2015). In this study, we explore the potential of a field spectroscopy technology as the basis of a precise, high-throughput, non-destructive tool for on-site determination of the concentration of algal proteins.

Measuring algae protein concentration and its associated phenotyping (Burnett et al., 2021) is critical for food industries that operate under strict health and environmental regulation and safety requirements (Christaki et al., 2015). However, algal phenotypes are highly diversified as a result of the complex interaction between algal physiology and biochemistry and environmental conditions (Großkinsky et al., 2015). Diversification can alter traits of interest, including quality and quantity. It is thus not surprising that very little is known about the dynamic between algal pigmentation and the concentrations of the associated proteins. Similarly, the recent literature is largely devoid of comprehensive spectral interpretations of the molecular composition of seaweeds and the spatial and temporal dynamics of their phenotypic expression as they interact with environmental stressors. Because the correlation between the traits of interest and VIS-NIR spectra is not linear, especially with regard to the protein fraction where absorbance wavelengths can overlap, linear tools may not accurately predict the content of active constituents, such as proteins, and more powerful interpretation techniques are needed (Yang et al., 2021). For this purpose, high-throughput, non-destructive analytical methods and phenotyping tools are being adopted to determine the algal growth rate for purposes of breeding and selection, and also for the study of algal physiology, biocomposition and nutrition characteristics, (Cozzolino, 2020) where crop phenotyping may be defined as the comprehensive assessment of complex plant traits and the basic measurements of individual quantitative parameters (Li et al., 2019). To date, crop phenotyping has been widely used in agriculture for determining the characteristics of leaves, fruits, and roots. Recently, it has found application in macroalgae for the selection of species for particular end-uses and for carbohydrate detection (Shefer et al., 2017) as well as for quantitative analysis of polyphenols and alginates (Beratto et al., 2017).

Important tools in crop phenotyping are imaging techniques that provide quantitative measurements characterizing the phenotype and biophysical crop parameters by utilizing the interaction between light and biomass reflectance, transmittance, and absorbance wavelength properties (Brook et al. 2020). Methods for collecting imagery data include remote sensing via commercial satellites and/or tools mounted on aircraft, which can yield different sections of the electromagnetic spectrum. However, for all these methodologies there is a tradeoff between the temporal resolution and the spatial resolution. Remote sensing for crop phenotyping that use multispectral and hyperspectral tools, are usually applied for determining nutrient status, growth rate, yield, and biomass mapping (Ohana-Levi et al., 2019; Paz-Kagan et al., 2020). For instance, the remote-sensing spectral tools developed for crop measurements have been applied for the management of forest diversity (Asner and Martin, 2016), and for crop management through determining crop nitrogen status (Elvanidi et al., 2018; Polinova et al., 2018), moderating field variability, or managing nitrogen fertilization (Ohana-Levi et al., 2019; Paz-Kagan et al., 2020). Remote sensing tools based on near-infrared reflectance spectroscopy (NIRS) have been broadly adopted by the agricultural and industrial sectors (Foley et al., 1998) as a means to rapidly identify important traits of interest in order to improve nutritive values. In that sense, diffuse reflectance is the internal reflected light attenuated by absorption of specific wavelengths, that scatters backward to external thallus and so constitutes a measurable expression of the phenotype pigmentation (Duppeti et al., 2017). The remote-sensing tools used in agriculture exploit the absorbance of particular wavelengths by specific functional groups as the biomass is exposed to light within the visible and near-infrared range (VIS-NIR; 400-1000 nm) (Yang et al., 2021). A few examples of the use of these tools may be cited. Beć et al. (2020) recognized the advantage of deep sample penetration by NIR radiation for in vivo examination in comparison to IR or Raman spectroscopy. Some studies demonstrated a correlation between light attenuation in the presence of chromophores and biomass traits associated with the nature of the C-N ratio and status in leaves (Ely et al., 2019). However, that study did not examine the correlation of absorption area with specific traits concentration. Others used this approach in microalgae for the rapid quantification of biomass density and pigments (Duppeti et al., 2017) and carbohydrates and lipids (Laurens and Wolfrum, 2013; Brown et al., 2014). In brown and red algae, most prior work on the mapping of proteinic traits used vibrational spectroscopy, such as Fourier transform infrared (FT-IR), in combination with analytical methods to identify protein-bound polysaccharides (Sumayya and Murugan, 2017; Beratto-Ramos et al., 2020) and antioxidant capacity (Beratto et al., 2017). These laboratory-based methodologies for testing of structural and functional properties of proteins necessitated the use of either dried, granulated, or powdered samples or samples recovered from precipitation (Abdollahi et al., 2019) or enzyme extraction (Vásquez et al., 2019).

The chromophore that is often used for categorization of the seaweed phylum (Olmedo-Masat et al., 2020), is part of a molecular complex involved in energy harvesting from sunlight for photosynthesis (Dumay et al., 2014). Phycobiliproteins (PBS) are a family of light-harvesting pigment-protein complexes (including phycoerythrin, phycocyanin and allophycocyanin) that are found in red seaweeds (Rhodophyta) and cyanobacteria, and are particularly efficient at harvesting light under low irradiance (Kannaujiya et al., 2017). PBS may constitute as much as 20% of the red seaweed dry matter and 50% of the seaweed water-soluble proteins. The survival of red seaweeds in diverse marine habitats ranging from deep water under low light irradiation to intertidal zones provides evidence for the efficient absorption of solar radiation and the high efficiency of energy transfer in these systems (Xie et al., 2021). However, the vast phenotype variability between and within groups of seaweeds that are affected by a complex of biotic and abiotic parameters poses an analytical challenge whose complexity requires new analytical tools. The use of spectral features and in-situ non-invasive methodologies for the identification of functional groups, such as proteins, has not yet been reported for seaweeds. Previous studies have indicated that the way to addressing this challenge lies in the application of VIS-NIR spectroscopy and neural networks and machine learning techniques to yield quantitative predictions of independent traits of organic material (Xu et al., 2019; Arinichev et al., 2021). The field spectroscopy approach measures point-by-point spectral radiance using a portable spectrometer. The main advantages of this tool are its low cost, high temporal and spatial resolution, and wide wavelength range across the VIS-NIR wavelengths (Polinova et al., 2018).

In this study, we thus developed and trained a novel neural network model to obtain, by means of a machine learning algorithm, predictions of protein content from phenotypic spectral features of the red seaweed *Gracilaria* sp. The seaweed was cultivated under six treatment regimes in a coastal setting in Haifa, Israel. *Gracilaria* sp. is known for its high level of essential amino acids, rapid growth rate, and commercial value. In 2018, it accounted for 10.7% of the world production of seaweeds (Chopin and Tacon, 2020). Its commercial value derives from its use as a source of agar (Armisen, 1995; Souza et al., 2012) and sulfated polysaccharides, which are used in pharmaceutical and biotechnology industries (Kazir et al., 2019). *Gracilaria* sp. was thus used as a model species for achieving the broader objective of this study—first to establish an accurate cultivation protocol for protein content accumulation in algae, and then to lay down the scientific and technological framework for predicting protein yield and moderating fluctuations in seaweed production.

## 2. Materials and Methods

### 2.1 Seaweed material, cultivation protocol and experimental design

The experimental setup was operated in two distinct seasons, summer (July) and fall (December) in 2020 (Fig. 1) in a seaweed tank culture system at the Israel Oceanographic & Limnological Research (IOLR) in Haifa, Israel (N 34°57’19’’ E 32°33’49’’). The system was composed of 18 40-L white fiberglass tanks which were filled with seawater and supplied with continuous aeration (Ashkenazi et al., 2020). About 100 g Fresh Weight (FW) of *Gracilaria* sp. were placed in each tank, and the algae were grown for 3-4 weeks under six treatments (n=3 tanks for each treatment protocol). The rationale for this experimental design was to explore possible alterations in protein content and seaweed phenotype as a result of a abiotic changes. The treatments included nutrient supplementation with two levels of nitrogen (added as NH_4_Cl) and phosphate (added as NaH_2_PO_4_). Moderate level additions comprised 1.0 millimole NH_4_ and 0.1 millimole PO_4_^3-^, and are designated (MN). High level additions comprised 2.0 millimole NH_4_ and 0.2 millimole PO_4_^3-^ and are designated (H). An additional set of six tanks containing as-received seawater (without ammonium or phosphate supplementation) were used as control (CON). According to Kress et al. (2014), the maximum nitrate and phosphate concentration in the Levantine Basin (Eastern Mediterranean Sea) is NO_3_^-^ ∼ 5.5 μmol kg^−1^, PO_4_ ^3-^ ∼ 0.21 μmol kg^−1^ respectively. Nitrogen and phosphate feedings were conducted every 4 days, and after each nutrient addition, seawater inflow was stopped for 24 h to allow maximal uptake by the seaweeds. To reduce excessive sunlight, nine tanks (three of each of the above mentioned groups, MN, H and CON, that were assigned to the different nutritional concentrations) were covered with two layers of a 5 mm mesh nets (hereafter designated low light; LI: A1, B1, C1: n=9) and nine others (again, three of each of the above mentioned nutritional concentration groups, MN, H and CON) with one layer (hereafter designated moderate light; MI : A2, B2, C2: n=9), yielding cutoffs of 75% and 50% of the incident sunlight, respectively. (See Fig. 1)

**Fig 1.**
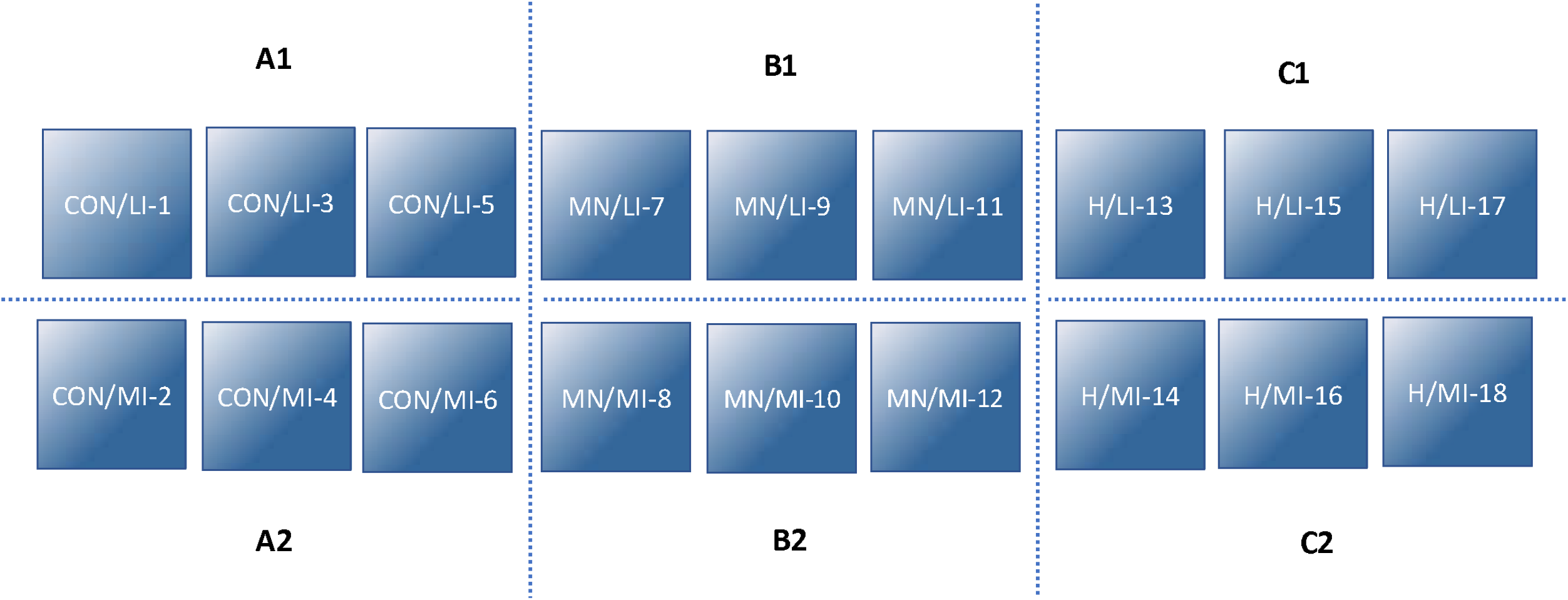
Schematic cultivation of Gracilaria sp. using treatments in a land-based outdoor setup. A1, A2 control group (CON); B1, B2 Moderate nutrients-enriched (MN); C1, C2 High nutrients-enriched (H). Biomass cultivated under low incident light intensity [(LI), n=9] and moderate incident light intensity [(MI), n=9].

Light intensity and seawater temperature were measured continuously by placing HOBO® Pendant® data loggers (ONSET UA-002-64, USA) in the tanks. Average light intensities during the summer were 298 and 155 μmol m^−2^ s^−1^ (depending on the light stress procedure), and 44 and 25 μmol m^−2^ s^−1^ during the fall. The average seawater temperature was around 28.5 °C during the summer, for all pools with no differentiation between trails. While during the fall season the average seawater temperature was 16.2 to 18.5 °C depending on one or two mesh net layers covering respectively. The biomass from each tank was harvested and weighed on a weekly basis for growth rate determinations. About 100 g FW of biomass were returned to the tanks. Algal chromophore reflectance properties from each tank were measured on a weekly basis with a field-portable spectrometer (Ocean Optics USB4000), acquiring data from seaweed thallus across the visible and near-infrared (VIS-NIR) range (400 to 1000 nm, with a resolution of 0.5 nm and an accuracy of 1 nm). The measured samples were retained for protein assessment using CHNS elemental analysis.

### 2.2 Seaweed growth analysis

The seaweed growth rate was determined on a weekly basis. After every harvest, the fresh seaweed biomass was drained of excess seawater and weighed, and specific growth rates (SGR, %FW per day) were calculated according to Neori et al., (2000), using the following formula:

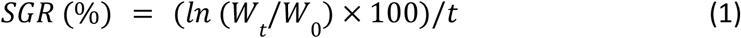

where W_0_ (in grams) is the initial wet weight of biomass and W_t_ (in grams) is the weight of biomass after t culture days.

### 2.3 In-situ spectral measurements of fresh seaweed thallus

To study the relation between protein concentration and seaweed phenotype, diffuse reflectance measurements were taken in-situ on a weekly basis in both seasons using a portable spectrometer. Randomly selected biomass thallus samples (FW) were taken out of the water from each of the 18 containers, 24 h after nutrient supplementation. These samples were measured in the visible and near-infrared (VIS-NIR) region of the electromagnetic spectrum (400-1000 nm, Fig. 2) on a fully sunny day. The spectrometer was calibrated according to Jackson et al. (1992) against a white Spectralon plate (Labsphere Inc., North Sutton, NH, USA). Bare fiber optic with 25° field-of-view positions was used for repeated collection of detailed spectra (Polinova et al., 2018). Reflectance measurements were conducted on various thallus areas that were usually cylindrical, blade-shaped branches. About 144 seaweed specimens were spectrally measured during the summer and fall trials, with 10-15 spectra repetitions for each sample.

**Fig. 2.**
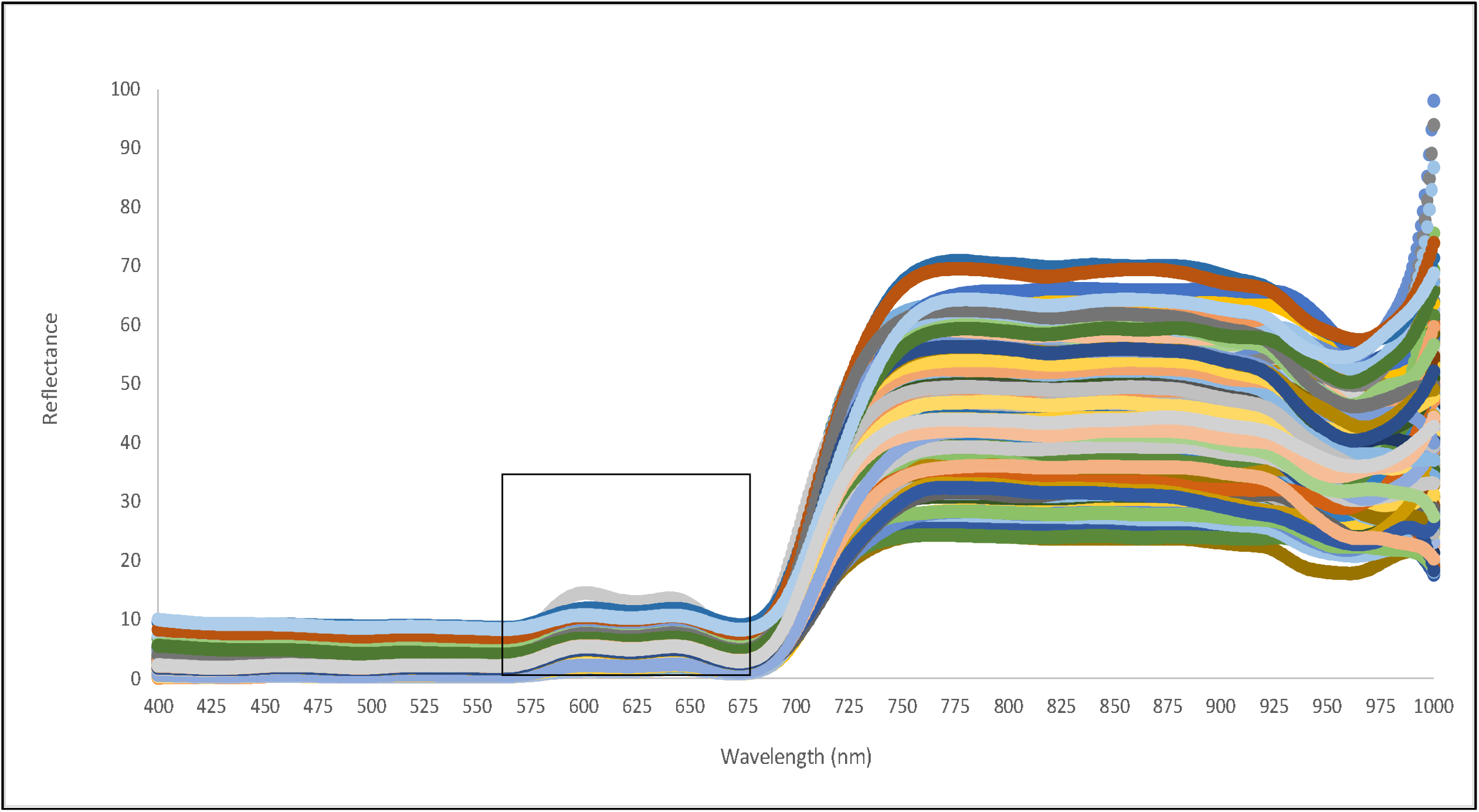
A representative reflectance spectrum of *Gracilaria* sp. measured in-situ on wet thalli. Measurements were conducted in the VIS-NIR region, 400–1000 nm. Absorbance at 560-680 nm was used to determine chromophores (phycobiliproteins) and protein components.

### 2.4 Elemental analysis and determination of seaweed protein content

Following spectral measurements, fresh seaweed specimens were taken on a weekly basis to the laboratory and stored in sealed plastic bags at -70 °C. Prior to elemental analyses, seaweed thalli were rinsed with tap water to remove salt, dried at 60 °C for 48 h, and milled to a fine particle size using a coffee grinder. The nitrogen content of the dried samples was measured using a Flash 2000 Organic elemental analyzer (Thermo Scientific, USA). About 2-3 mg of the dried and milled material from each specimen (n=144) were combusted at 960 °C according to manufacturer’s protocol. The nitrogen content of the dried samples was used to determine protein content, assuming a nitrogen-to-protein (N-prot) conversion factor of 5.0, which has been shown to be appropriate for marine seaweeds (Angell et al., 2016).

### 2.5 Spectral wavelength selection and data pre-processing

Spectral measurements were obtained for phycobiliproteins, which are the main light-harvesting proteinic pigments in Rhodophyta. For these chromophores, namely, pink/purple-colored phycoerythrin (PE), blue colored phycocyanin (PC), and bluish-green colored allophycocyanin (APC) (see Fig. 3), light energy is transmitted from those absorbing at green wavelengths to those absorbing at red wavelengths (Dumay et al., 2014; Dumay and Morançais, 2016; Kannaujiya et al., 2017). In our case absorption was observed mainly in the wavelength range 560-680 nm. To overcome the low selectivity of VIS-NIR spectral information due to the texture, size, and geometry of the seaweed thallus, we used the Kubelka-Munk model for the diffuse reflection scattering coefficient, using the following equation which is an analog to absorbance transformation in transmission (Ma et al., 1987):

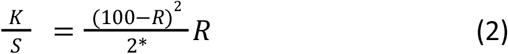

where reflectance R of seaweed thallus pigment is related to the ratio of the absorption coefficient of seaweed chromophore K and the scattering coefficient S (Militký 2011). The ratio of the coefficients is proportional to the pigment concentration (Ma et al., 1987). Pre-processing of spectral data that used as an input to the Artificial Neural Network (ANN) also included identification of minimum values, baseline correction to smooth and normalize the scatter spectra measurements (Engel et al. 2013; Cozzolino 2020), and absorption depth calculation. To further improve the model, as a first step spectral measurements repetitions of each specimen were averaged. However, due to variations in the chromophore phenotype in the samples derived from seaweed subjected to different environmental stressors (nutrient supplementation and subtraction of incident light) and the spectral expression of these fluctuations in the same measured sample, all spectral measurements were considered.

**Fig. 3.**
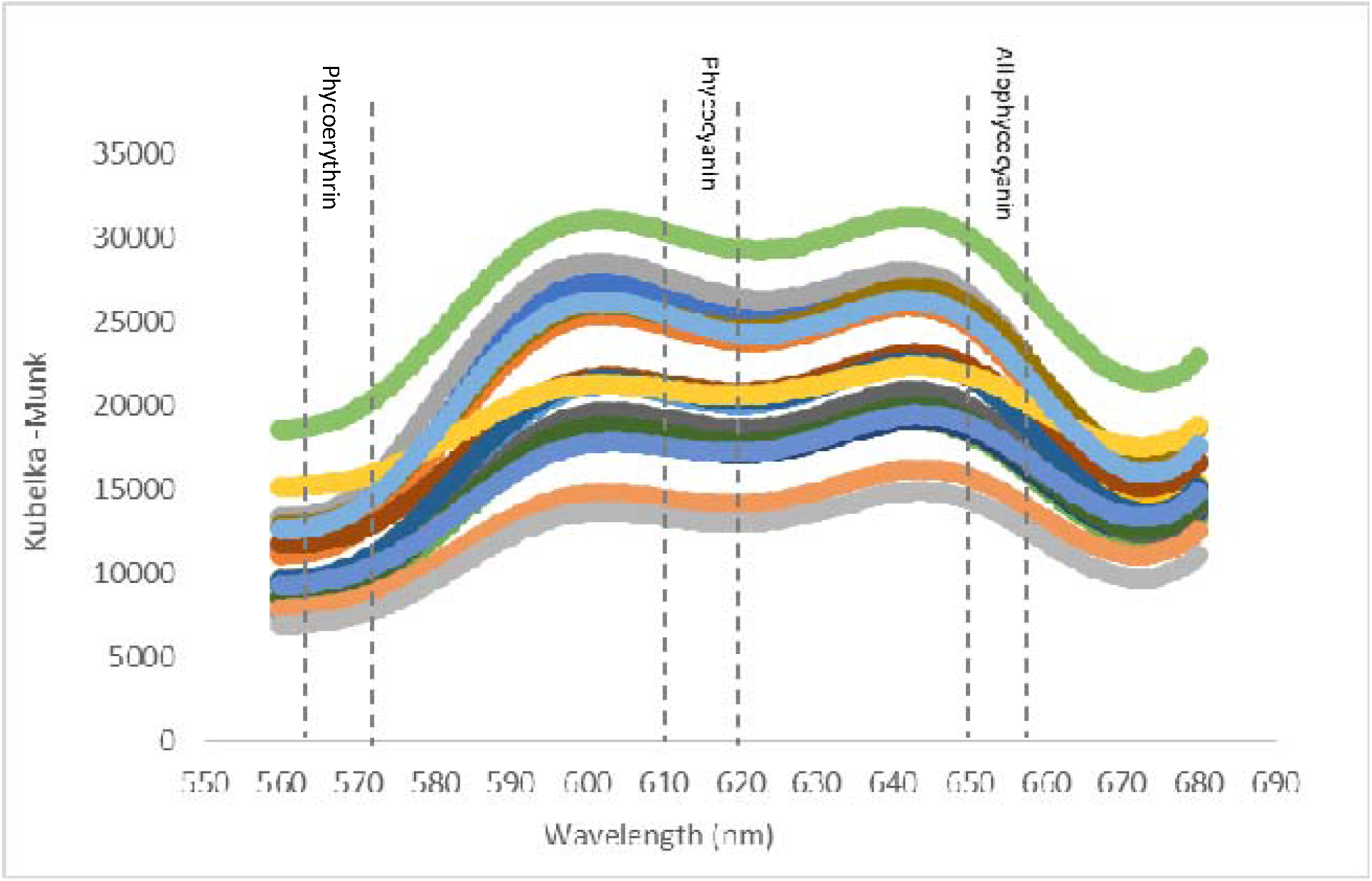
Phycobiliprotein absorption area. Kubelka-Munk normalized absorbance features of phycobiliproteins (PBS) in *Gracilaria* sp. PE: λ_max_ = 560-570 nm; PC_max_ = 610-620 nm; APC_max_ = 650-655 nm. These antenna pigments are covalently amino acids (Aryee et al., 2018) enabling light energy to transfer through this pathway: *PE* →*PC* →*APC* →*chl a* (Dumay and Morançais, 2016).

### 2.6 Data processing

A backpropagation artificial neural network (BP-ANN) that has a solid theoretical basis and is widely used in various fields (Huang and Foo, 2002), was applied to process the spectral data. The BP algorithm is, in general, the most common learning algorithm used to construct nonlinear models. BP-ANN is a multilayered network based on the generalized Woodward-Hoffmann learning rule and weight-trained differentiable nonlinear functions with a strong learning ability. The BP-ANN method was improved by introducing the momentum factor and the weight control algorithm (MBP-ANN). The MBP-ANN was conducted on the MATLAB platform. BP-ANN model-building and precision validation were implemented using the Neural Network Toolbox. The structure of the MBP-ANN was composed of three layers: input, hidden, and output layer. The pre-processed spectral data (as described in Section 2.5) was used as the input to MBP-ANN for the development of a protein content prediction algorithm. In estimating the dependency function (Polinova et al., 2019), we found a non-linear correlation between seaweed specimen reflectance properties and the measurements of protein content for the same specimen obtained from nitrogen elemental analysis (converted to protein content by N-prot factorial multiplication). An input layer with pigment intensity per wavelength was propagated from input to output through neurons, random weights and bias, and trained, validated, and tested. The input of one layer consisting of 10 neurons of the red wavelength absorbance spectral area (670-680 nm) was broadened to 114 neurons in 1-nm intervals to include the absorbance area of the chromophores (560-674 nm). Training methods included backpropagation to fine tune the weights and bias of the model parameters and the Bayesian regularization technique (Foresee and Hagan, 1997) for better generalization. Since the weight and bias of the model, which vary depending on the hyperparameter values, can affect the model performance, the model performance was analyzed by changing the model structure and hyperparameters to ensure optimal outcomes. Therefore, the performance of the initial MBP-ANN model was optimized by changing its hyperparameters, and structural Bayesian optimization was employed to determine the optimal combination of hyperparameters to enhance the prediction performance. Based on the Bayes Theorem, a surrogate model, which is a probabilistic model of the objective function, searches for specific hyperparameters yielding the maximum or minimum performance. A rectified linear unit (ReLU) propagated backward computed errors to update the model parameters, and then the predicted normalized means were compared to the actual protein content results obtained from the CHNS elemental analytical analysis.

MBP-ANN training, testing, and validation of the seaweed thallus phenotype were conducted on data obtained from the field measurements for the two seasons, normalized and processed using the Kubelka-Munk equation (Eq. 3) with 1000 iterations, until fully connected.

The loss function used for model optimization to reduce overfitting and large weights was given by the following equation:

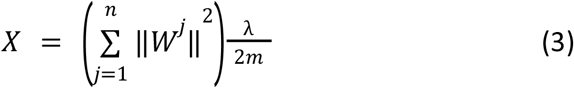

where *n* - the number of layers, *W* - the weight matrix for the j layer; *m* - the number of inputs and λ - the regularization parameter.

The input data set for the MBP-ANN consisted of 164,160 spectral measurements of which 70% were used for training, 15% for testing, and 15% for validation. The output data was the prediction of protein content (% DW) that fully connected with pigment intensity per wavelength within the VIS-NIR area (560-674 nm).

### 2.7 Post-processing

To further validate the model, we performed an external validation trial for four consecutive weeks during the fall of 2021 for six cultivation regimes (including control). To smooth specimen variability due to abiotic stress, the trial setting was narrowed to six containers, and three replicates of Gracilaria. sp. were taken randomly from each container (i.e., 18 samples). To assess the ability of MBP-ANN to automatically extract useful patterns and predict protein content from a data set that was not previously used, Fifty-four samples of the cultivated fresh biomass were classified in-situ via reflectance spectral features measurements (560-680 nm) using absorption depth. The input data included field measurements on seaweed thallus phenotype, normalized and processed using the Kubelka-Munk equation. The predicted normalized means were compared to the actual protein content results obtained from N content elemental analysis of the dried samples, which were converted to protein content (% DW) using the same N-prot factor of 5.0. Model performances were assessed in terms of the regression coefficient R2, the mean square error (MSE) and the root mean square error (RMSE) for the validation results, according to the following equation:

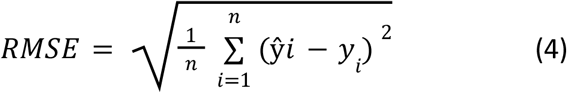

where *yi*, ŷ*i* - measured and predicted values of protein content, and *n* - the number of samples.

## 3. Results

### 3.1 Seaweed growth performance and protein content

A large variation in SGR was observed among and within groups of seaweed and between seasons (Fig. 4; Table 1). During the summer, SGR ranged between 3.55% and 10.45% per day, which was much higher than the *Gracilaria* sp. SGR during the fall (between 0.55% and 4.23% per day). However, in both seasons the lowest growth rate was observed for the control group (A2) under conditions of moderate light stress and no nutrient supplementation (3.55–7.0% per day during the summer and 0.55–2.41% per day during the fall). The maximum SGRs of 10.45% and 4.23% per day were observed for low light intensity and high nutrient enrichment (group of containers C1) during summer and fall, respectively. Analysis also revealed a large variation in protein concentration across seaweed samples. In general, during the fall, *Gracilaria* sp. exhibited much higher protein content consistently in all treatments in comparison to protein content measured during the summer (7.43–17.43 % DW; 1.56–6.05 % DW, respectively). As anticipated, nutrient supplementation and light stress contributed to protein accumulation in the seaweed biomass. The highest protein content was observed during the fall under exposure to moderate incident light and a high level of nutrient supplementation (17.43% DW, group C2). During the summer, the seaweed exhibited the highest protein content under exposure to low incident light and high nutrient enrichment (6.05 % DW, B1). No correlation was found between SGR and protein content. Therefore, SGR was not used as a proxy in ANN training.

**Table 1.** The range of specific growth rate and protein content

|  | SGR range (% / day)* |  | Protein content range (% DW)* |  |
| --- | --- | --- | --- | --- |
|  | Summer | Fall | Summer | Fall |
| A1 | 4.17 - 7.04 ± 1.41 | 1.16 - 2.96 ± 0.75 | 1.56 - 4.97 ± 0.16 | 8.31 - 9.15 ± 1.08 |
| A2 | 3.55 - 7.0 ± 1.74 | 0.55 - 2.41 ± 1.18 | 3.30 - 4.40 ± 0.16 | 7.43 - 8.27 ± 0.71 |
| B1 | 6.64 - 7.56 ± 1.57 | 3.16 - 3.85 ± 0.69 | 5.02 - 6.05 ± 0.26 | 12.79 - 13.64 ± 0.97 |
| B2 | 8.17 - 9.53 ± 1.97 | 1.80 - 2.82 ± 1.97 | 3.07 - 4.17 ± 0.15 | 11.62 - 14.16 ± 3.47 |
| C1 | 8.10 - 10.45 ± 1.26 | 2.29 - 4.23 ± 1.26 | 4.25 - 4.97 ± 0.28 | 14.57 - 15.56 ± 2.29 |
| C2 | 8.35 - 10.17 ± 1.50 | 2.21 - 2.55 ± 1.78 | 3.10 - 4.57 ± 0.10 | 17.07 - 17.43 ± 4.07 |

\* Numbers are in average terms
*Gracilaria*. sp. range of averages ± SD of specific growth rate (% /day) and protein content (% DW) during the summer and fall seasons of 2020, calculated for each treatment regime (A1-2; B1-2; C1-2) at four time sequences (T1-T4). The range describes fresh biomass specific growth rate (% day<sup>-1</sup> FW), and in percent of dried biomass for protein content determination as calculated from elemental analysis.

**Fig. 4.**
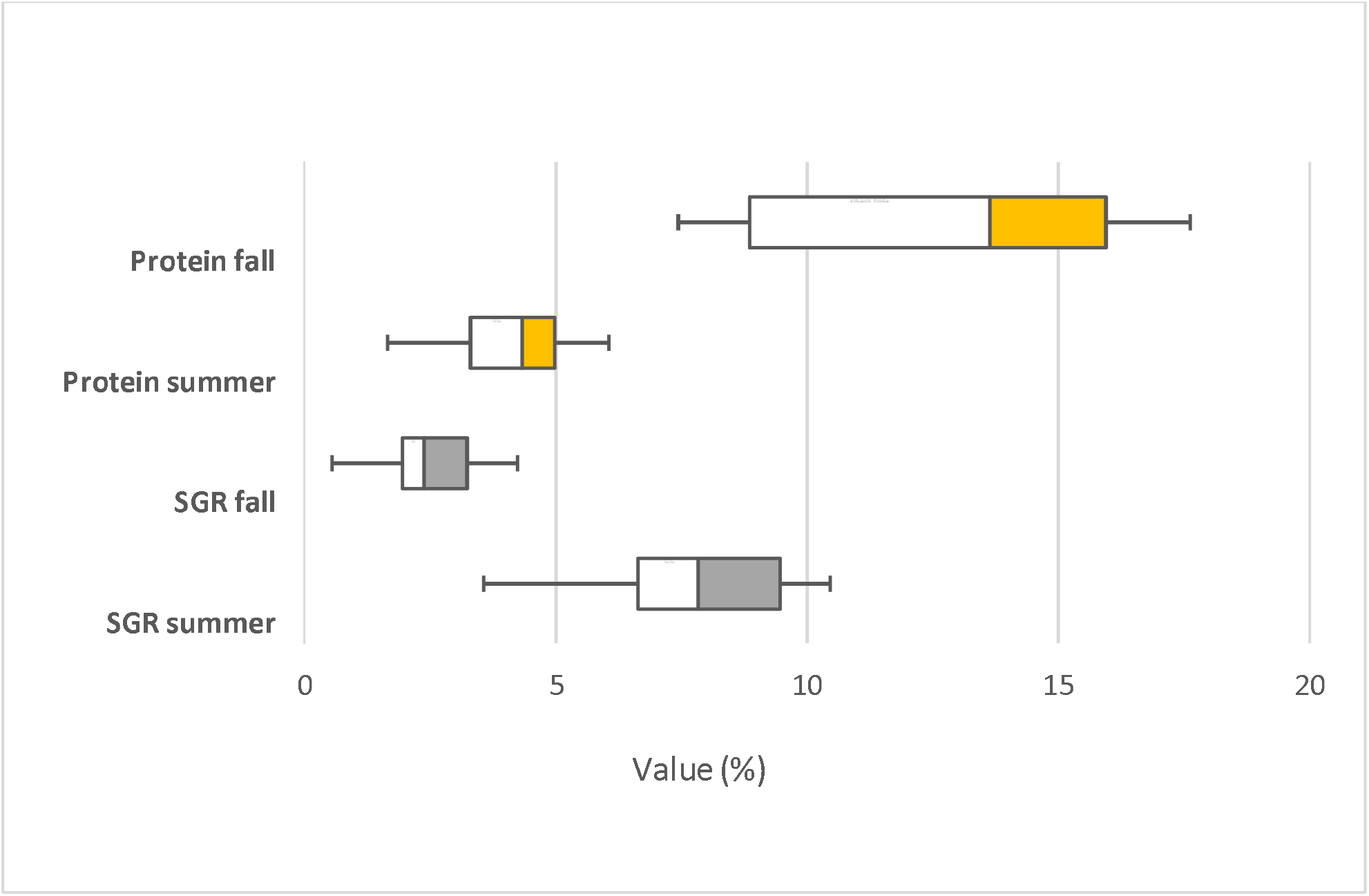
Growth rate and protein content of *Gracilaria* sp. cultivated in the summer (July) and fall (December) trials of 2020. Gray: Specific growth rate, by season [(SGR), % /day]; Yellow: Protein content (% DW). Boxplots bars represent the minimum and maximum observed values.

### 3.2 Algal thallus phenotype and ANN training results

Significant variability was observed in algal phenotypes in all containers throughout the trials, including at the specimen level. Thallus phenotype pigment ranged from light yellow and yellowish-brown to pink and dark red (Fig. 5).

**Fig. 5.**
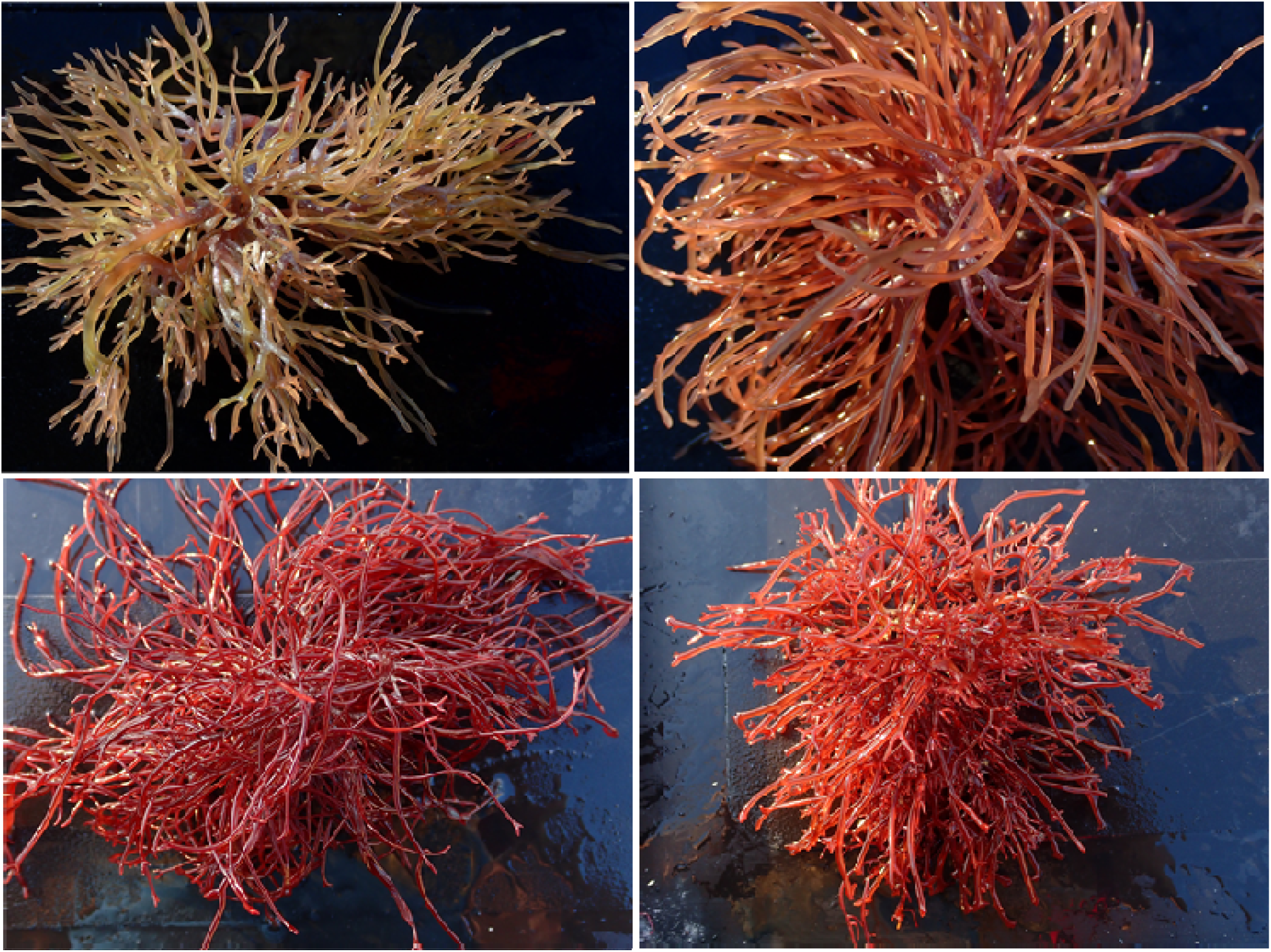
Cultivated Gracilaria sp. specimens. A large variability in phenotype and thallus colour were observed during the fall season, reflecting seaweed protein acclimation mechanism to changes in nutrient supply and incident light. Cultivated specimens were classified via reflectance spectral features across the phycobiliprotein absorption area (560-680 nm) using absorption depth.

The MBP-ANN trained with 1000 iterations to connect pigment intensity with protein absorption in a narrow zone of the red wavelength 670-680 nm resulted in relatively low prediction accuracy of R^2^ = 0.74. Broadening the absorption area to include chromophores at 560-674 nm for data training produced much higher values of the correlation coefficient (R^2^). The initial optimal value of mu was 0.9899, and the optimal number of hidden neurons, which are structural variables for the model, was 10 (Table 2). The optimized model performances for the training and test data were compared with the performance of the initial model (Table 3). The R^2^ for the test data was 0.95 (p < 0.01) for the optimized model (see also Fig. 6). Moreover, the RMSE for the optimized model was 0.84 indicating a higher accuracy than for the initial model (RMSE = 4.6, Table 3).

**Table 2.** Hyperparameters for MBP-ANN optimization

| Hyperparameter | Description | Optimized value |
| --- | --- | --- |
| Neuron size | Number of neurons in the hidden layer | 10 |
| mu | Initial mu setting | 0.9899 |
| mu decrease | mu decrease factor | 0.8692 |
| mu increase | mu increase factor | 6.8909 |
| Min gradient | Minimum performance gradient | 6.45E-06 |

**Table 3.** Performance analysis for the MBP-ANN prediction model

|  |  | R<sup>2</sup> | RMSE |
| --- | --- | --- | --- |
| Initial model | Training | 0.92 | 3.85 |
|  | Test | 0.89 | 4.6 |
| Optimized model | Training | 0.97 | 0.64 |
|  | Test | 0.95 | 0.84 |

Prediction accuracy for results of initial and optimized model. $R^2$ = coefficient determination, RMSE = root mean square error.

**Fig. 6.**
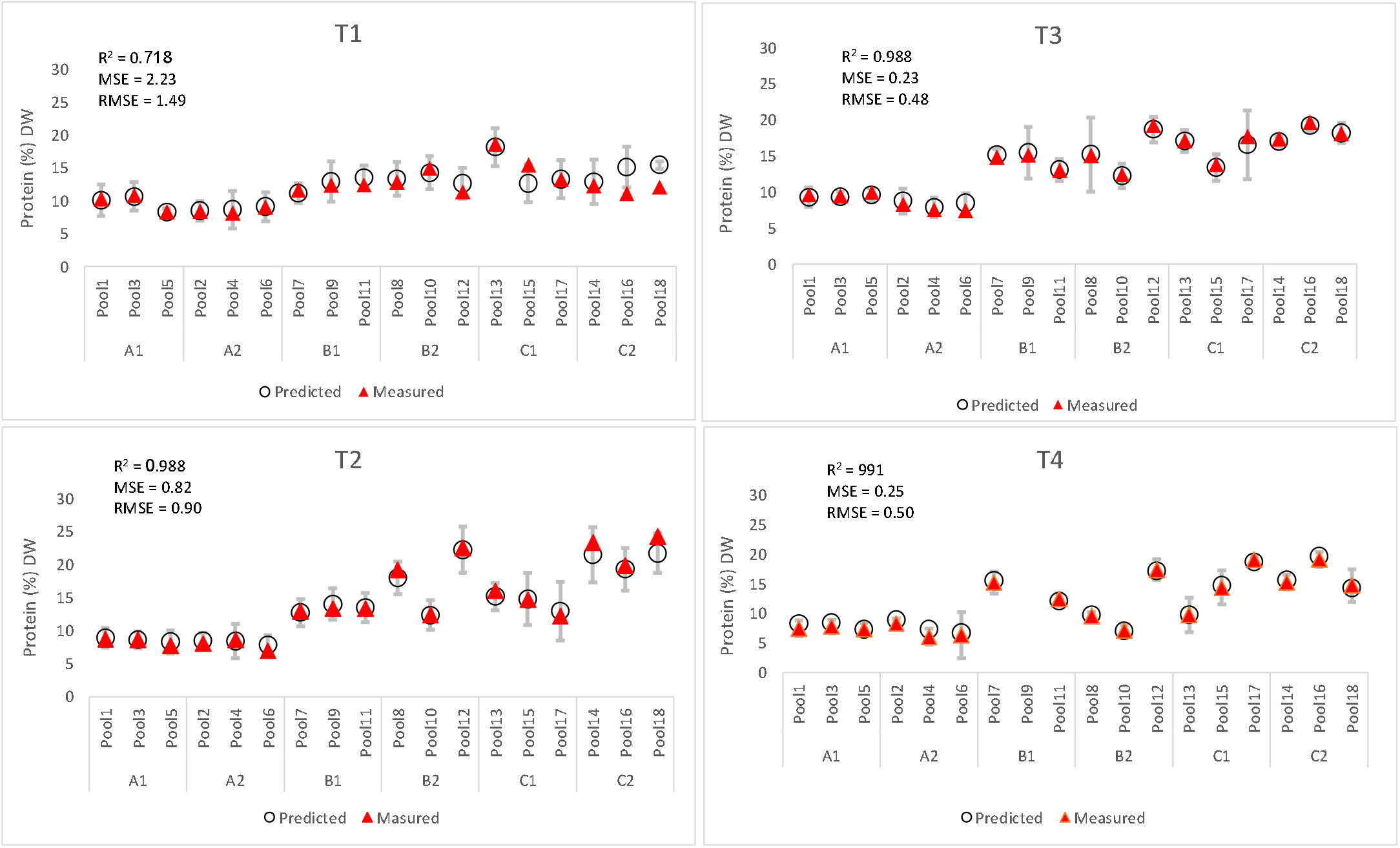
Model prediction performances. Mean ± standard error of predicted protein content values in the algae biomass (% DW) as obtained from the MBP-ANN model, in comparison with laboratory results obtained from the CHNS analytical analysis of N content converged to protein content (see Table 5). The graphs present measurements during four consecutive weeks (hereafter designated T1-T4) conducted during the fall season of 2020. Range of R^2^: 0.718-0.991; Range of MSE: 2.23-0.23; Range of RMSE: 1.49-0.48.

### 3.3 MBP-ANN external validation results

The robustness of the model was demonstrated by the highly accurate prediction of protein content (% DW), which used the input data of spectral features that were collected during the external validation trial in fall 2021, without any training iteration (see: Fig. 7; Table 4). The R^2^ value of the external validation trial was 0.916, the MSE was 0.24, and the RMSE equal to 0.44.

**Table 4.** Model performances for the subsequent experiment conducted during fall 2021.

| ANN prediction model performances |  |  |  |
| --- | --- | --- | --- |
| Protein content (% DW) | $R^2$ | MSE | RMSE |
| External validation | 0.993 | 0.247 | 0.452 |

The values represent protein contents predicted by MBP-ANN versus CHNS elemental analysis

**Table 5.** Elemental analysis of Gracilaria sp. samples from six treatment regimes

| *Gracilaria* (% DW) |  |  |  |  |  |  |
| --- | --- | --- | --- | --- | --- | --- |
| Treatment | C | H | N | S | C:N | Protein (N * 5) |
| 1 (CON/LI) | 28.31 ± 1.3 | 4.20 ± 0.16 | 0.90 ± 0.07 | 3.07 ± 0.25 | 31.74 ± 2.74 | 4.5 |
| 2 (CON/MI) | 27.94 ± 0.21 | 3.96 ± 0.03 | 0.63 ± 0.06 | 2.64 ± 0.36 | 44.80 ± 4.76 | 3.2 |
| 3 (MN/LI) | 27.08 ± 2.48 | 4.11 ± 0.37 | 2.86 ± 0.04 | 2.06 ± 0.19 | 9.44 ± 0.82 | 14.3 |
| 4 (MN/MI) | 26.94 ± 1.97 | 4.04 ± 0.27 | 2.82 ± 0.11 | 2.16 ± 0.09 | 9.54 ± 0.56 | 14.1 |
| 5 (H/LI) | 30.75 ± 0.35 | 5.0 ± 0.06 | 4.61 ± 0.05 | 2.08 ± 0.12 | 6.66 ± 0.07 | 23.1 |
| 6 (H/MI) | 27.03 ± 1.25 | 4.35 ± 0.16 | 4.01 ± 0.30 | 2.21 ± 0.18 | 6.78 ± 0.71 | 20.07 |

CHNS elemental analysis (%) of *Gracilaria* sp. obtained from six treatment regimes (CON = control; MN and H = Moderate and High nutrient enrichment; LI and MI = Low and Medium incident light exposure) on a dry basis (DW); C:N the ratio carbon to nitrogen and estimated protein content. Values are means ± SD.

**Fig. 7.**
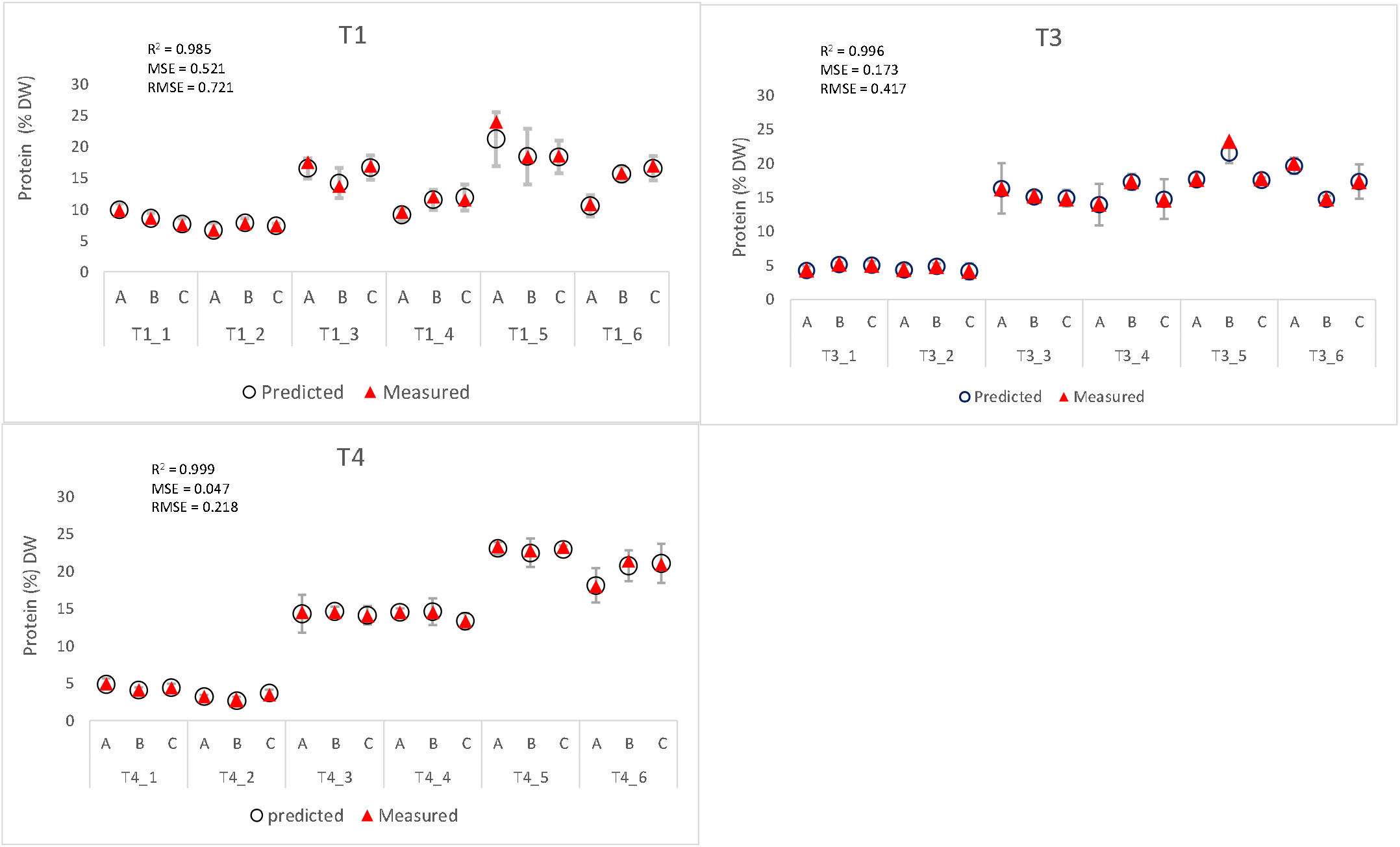
Accuracy of prediction for the external validation trial. Values of specimen protein content (% DW) in means ± standard error, under six cultivation regime per time sequence (T1, T3, T4), were obtained from the external validation trial during the fall season of 2021. The model exhibited mean R^2^ value of 0.916, MSE of 0.24, and RMSE of 0.44. The ANN model used the spectral features that were collected during this trial and generated a prediction without any training iteration.

## 4. Discussion

This paper evaluates for the first time the accuracy of the MBP-ANN algorithm in using in-situ-acquired diffuse reflectance measurements in the VIS-NIR spectrum to predict the protein content in the red seaweed *Gracilaria* sp. We found that when using data from wide spectral features of phycobiliproteins absorbance area (560-674 nm, R^2^=0.993) rather than the narrow spectral window of the red wavelength absorbance area at 670-780 nm (R^2^=0.74), the algal pigment phenotype is a reliable predictor of the analyzed nitrogen concentration. The high pigment phenotype diversity that was observed throughout the trials (Fig. 5; Fig. 8) is a manifestation of alterations in protein concentration (Barth, 2007) as changes occur in response to the environmental conditions. The pigment intensity of the algal thallus does not only change temporally for a given group of seaweeds and between treatment regimes, but also varies among different parts of the same specimen. For instance, the blade tissue has a higher pigment concentration than the branches, which manifest as differences in the wavelengths of unabsorbed diffuse reflectance scattering. Olmedo-Masat et al. (2020) also recognized pigment phenotype concentration differentiation across different parts of the brown seaweed *Macrocystis pyrifera*. It is evident that seasonality (Khairy and El-Shafay, 2013), incident light intensity, and nutrient enrichment affect protein accumulation in the algal biomass (Fig. 8). These findings confirm previous studies, in which the pigment phenotype confers on *Gracilaria* species, like other rhodophytes, an evolutionary advantage in adaptation to environments of low light intensity (Saluri et al., 2020; Xie et al., 2021) by transferring excitation light energy of PE to PC to APC (Grabowski and Gantt, 1978; Dumay and Morançais, 2016). This acclimation mechanism, which determines seaweed thallus pigmentation (Fig. 3; Fig. 5; Fig. 8), can be predicted (Duppeti et al., 2017) and therefore it can be partially controlled. However, phenotypic expression is not linearly correlated to environmental conditions. Furthermore, overlapping bands in the spectrum, which are also due to the complex three-dimensional composition of protein molecules, can hide spectral information relating to the concentration of a particular trait (Barth, 2007; Yang et al. 2021). As a consequence, popular multivariate regression techniques for the identification of absorption bands and their correlation with a trait of interest for content prediction, such as multiple linear regression, partial least squares regression (PLSR), principal component regression (Beratto et al., 2017; Duppeti et al. 2017; Ely et al., 2019) were not applicable in our study. We used the physical and biochemical features of the absorption depth in diffuse reflectance spectroscopy of the phycobiliproteins and the Kubelka-Munk equation as inputs to MBP-ANN and ML algorithms. These algorithms were able to generate highly accurate semi-quantitative prediction of protein content through an iterative process in which pigment intensity as a function of the wavelength within the VIS-NIR area (560-674 nm) and protein concentration (% DW) were fully connected. The performance accuracy of the model (see Table 4) also served to validate the reliability and accuracy of the N-prot conversion factor of 5.0 (Angell et al., 2016) as a quantitative method to determine the protein content of red seaweeds (Table 5). Since not all the nitrogen is contained in proteins, and other nitrogen-containing molecules are present in the algae (such as inorganic nitrogen and free amino acids), previous studies that suggested average conversion factors of 4.59 (Lourenço et al., 2002; Kazir et al. 2019) 4.92 (Mæhre et al. 2014), or 6.25 (Tabarsa et al. 2012) for red seaweeds would lead to under- or over-estimation of protein content.

**Fig. 8.**
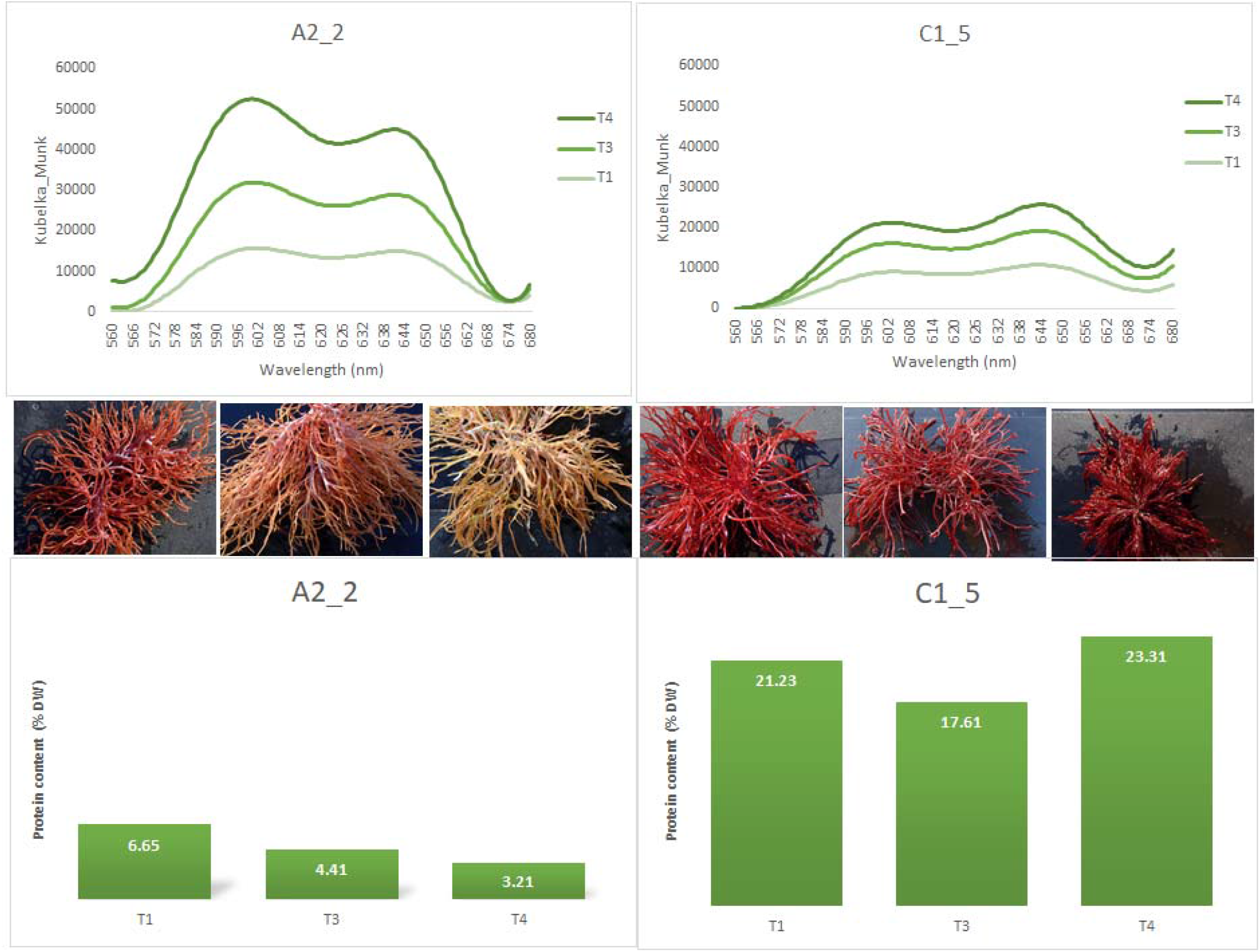
Spectral, chemical and phenotyping features used for model development. Normalized Kubelka-Munk spectral features for *Gracilaria* sp. for experimental treatment regimes showing minimum (A2-2: Control group: no nutrient addition, moderate *incidGraent* light) and maximum (C1-5: high nutrient addition, low incident light) protein accumulation (% DW) through three weekly time sequences (T1, T3, T4) *. The measured specimens were photographed at each time sequence. *Weather conditions prevented spectral measurements during the second time sequence (T2) of the fall season, 2021

### 4.1 Growth rate versus protein accumulation

In most of the previous studies that sought to assess the prevalence of traits of interest in seaweed and their physiological reaction to the influence of environmental factors, the daily growth rate of biomass was also addressed (Pliego-Cortés et al., 2017; Shefer et al., 2017; Bermejo et al., 2020; Zepeda et al., 2020) but without assessing the effect of the growth rate on the accumulation of the desired trait. Saluri et al., (2020), for instance, found a weak correlation between biomass density and yields of phycobiliproteins. Our study showed that SGR was affected by seasonality (Fig. 4), and culture conditions and duration. A comparison between SGR and protein concentration (Fig. 9) did not reveal any significant patterns. Furthermore, growth rates changed for individual specimens between and within treatments during cultivation periods, but no correlation was found to protein content. While SGR measurement is an important factor for yield prediction of fresh biomass, it was an insignificant factor for prediction of protein content and was therefore not used as an input to the MBP-ANN and ML algorithm.

**Fig. 9.**
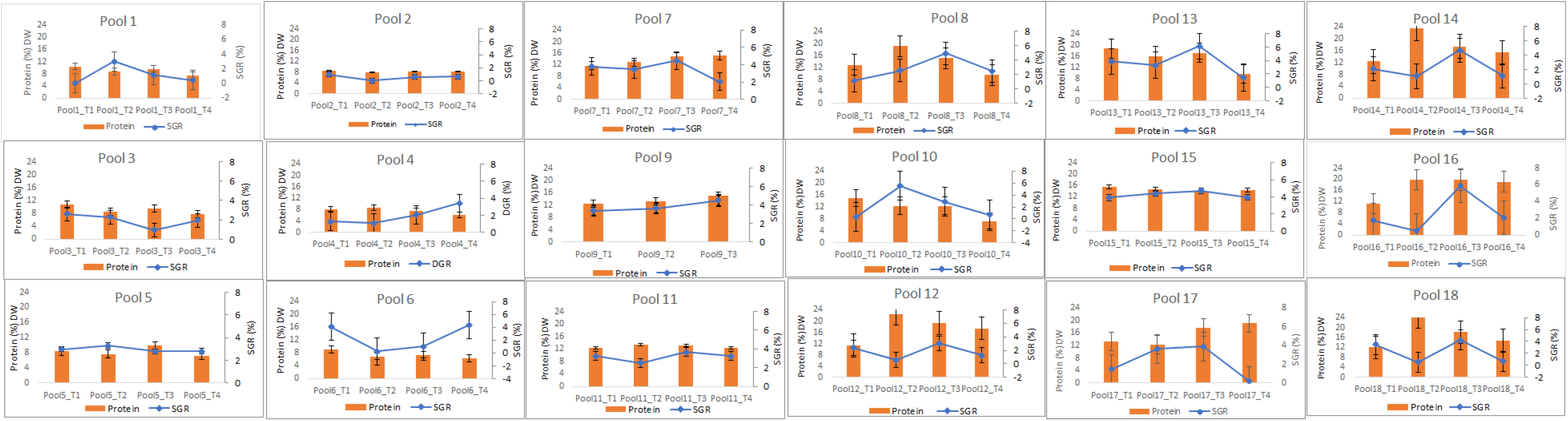
A comparison of seaweed growth rate and protein content. Specific growth rate (SGR) measured as FW (% /day) over four consecutive weeks in comparison with protein content (% DW) for the cultivated *Glacilaria* sp. during fall season. Data was analyzed per specimen per container (n=18) and figures are in average ± SD (error bars)

### 4.2 Reduction of variability by manipulation of cultivation regime

Food industries operate under strict health and environmental regulations and safety requirements. Therefore, knowledge of protein concentration and the associated phenotype is critical, since phenotype diversity of pigment intensity is actually an expression of protein trait alterations. This requirement underlies the goals of this study to identify (by means of diffuse reflectance spectroscopy and use of the ML algorithm) optimized cultivation conditions for protein content enhancement and to moderate fluctuations. The outdoor cultivation setting, composed of 18 containers under six cultivation regimes during the trial, was reduced to six containers during the external validation trial, during which three specimens replicates were taken from each of the six containers. The highest variation in protein content, as shown by the coefficient of variation (CV) per treatment regime during the training trial was 47% of the laboratory analysis results and 37.87% of the results predicted by the algorithm (Fig. 10). This variability was observed under moderate light reduction and moderate nutrient enrichment (B2). A substantial reduction in CV was achieved in the external validation trial, where the greatest variation was 18.45% (measured) and 18.55% (predicted) under a low level of incident light exposure and a high level of nutrient enrichment (C2). The more unified cultivation conditions, the smaller variations observed in the spectral features and specimen phenotype.

**Fig. 10.**
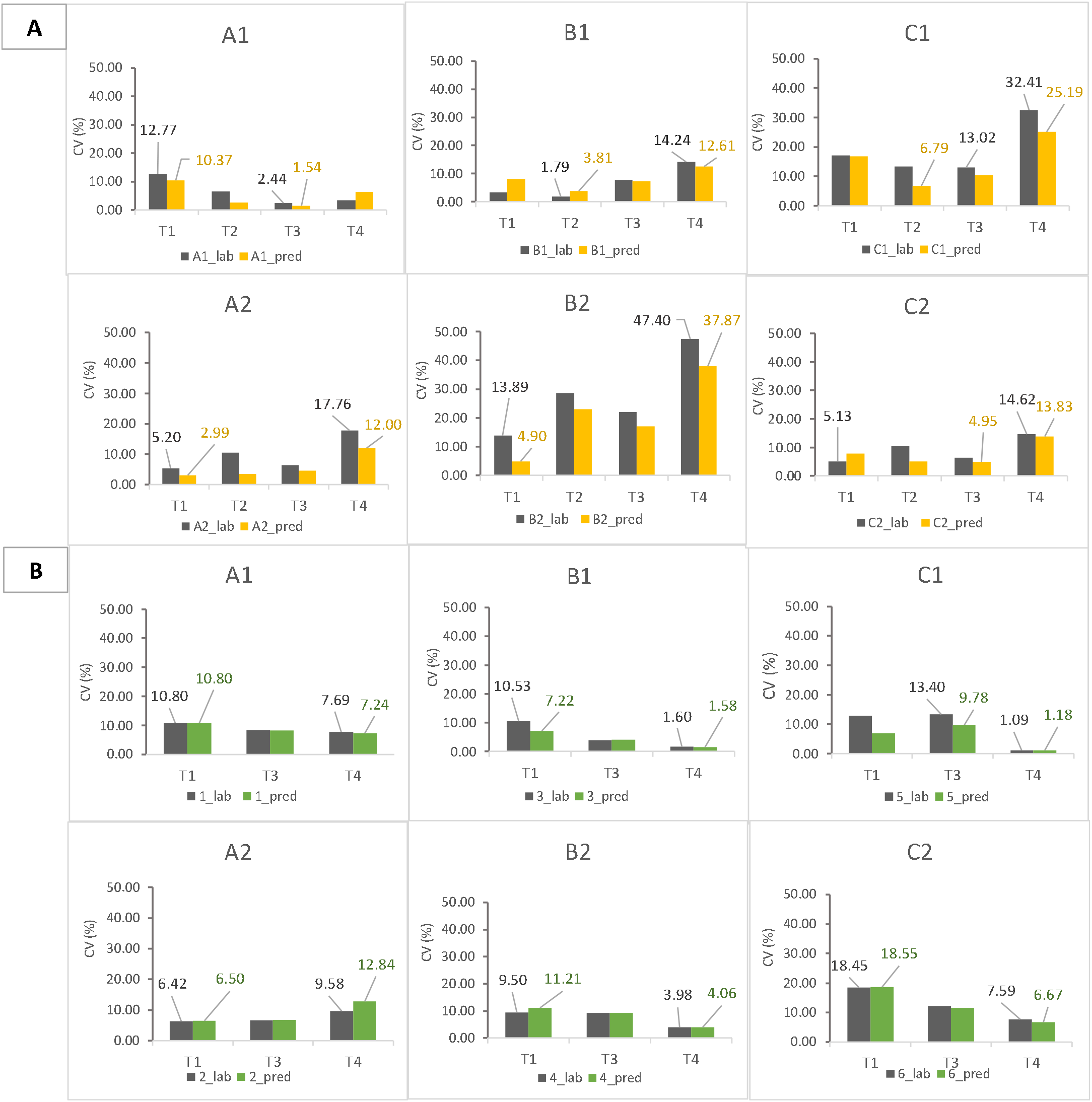
Range of coefficient of variation (CV). CV of protein content in *Glacilaria*. sp. per cultivation regime (A1-2, B1-2, C1-2) per time sequence (T1-T4) at two trail sessions: training (A) and external validation (B). Grey columns are representing measured results of lab analysis and colored columns are representing ANN predicted results. Numbers above columns representing minimum and maximum figures per treatment regime (A1-2; B1-2; C1-2).

### 4.3 Limitations and future directions

In this study, the generalized MBP-ANN model produced highly accurate predictions of protein content from highly diversified spectral features of *Gracilaria* sp. However, it may be necessary to adjust both prediction accuracy of protein content and cultivation protocols before they can be applied to other algal species and cultivation infrastructures. In this context, additional work on the characterization of changes in the protein concentration and its phenotype expression caused by specific cultivation parameters could result in better estimates for the weights of environmental parameters used for stress initiation in accordance with specific desired outcomes and yield improvement.

## 5. Conclusion

MBP-ANN is a powerful tool that successfully extracted useful patterns from a non-linear dataset of diffused reflectance seaweed phenotype features and performed highly accurate protein content prediction in a species of red seaweed without relying on chemical analysis. The accuracy of the model was validated in an external validation trial, which improved the prediction correlation coefficient performance (from R^2^ = 0.921 to R^2^ = 0.993). Fine-tuning of the weights and bias of the model parameters during the training and the Bayesian regularization technique for better generalization contributed to higher accuracy predictions of protein content, despite the high diversification of the pigment color. The phycobiliproteins absorption area (560-674 nm) was found to be much more informative than the narrow spectral wavelengths of the red band only (670-680 nm; R^2^ = 0.74). The non-destructive, high-throughput model developed here can support decision making for improvement of protein yield.

Furthermore, the findings of this study can inform future research exploring the potential of algae to become an important source of highly nutritive protein. We have shown that the seaweed pigment phenotype is a reliable indicator for protein concentration, which can be accurately predicted and managed using non-destructive high-throughput capabilities, thus enabling decision-making in-situ. These capabilities can foster precision in production, facilitate efficient use of resources, aid in compliance with regulations, reduce risks and, in some cases, prevent economic loss. The study has also established the analytical and technological foundations for a more generic model, in which different traits can be identified and quantified in-situ for better exploitation of different algal components.

## Abbreviations

ANN: artificial neural network
APC: allophycocyanin
BP: backpropagation
CV: coefficient of variation
DW: dry weight
FTIR: Fourier transform infrared
FW: fresh weight
IR: infrared
ML: machine learning
Mu: training gain in ANN
MBP: momentum factor and weight control algorithm
N-prot: nitrogen-to-protein
NIR: near infrared
PBS: phycobiliproteins
PC: phycocyanin
PE: phycoerythrin
SGR: specific growth rate
VIS-NIR: visible near infrared

## Acknowledgments

We thank the Department of Natural Resources and Environmental Management and the Petrie Foundation for financial support, and the Israel Oceanographic & Limnological Research for use of cultivation infrastructure facilities. We also thank Dr. Rachel Edrei the manager of the chemical and surface analysis laboratory at the Faculty of Chemistry, Technion (IL) for providing elemental analysis infrastructure and professional support. We thank Dr. Roland Mureinik from Ben-Gurion University (IL) for academic and language editing services.

## Author’s contributions

All authors contributed to writing the manuscript: NTS and AB for the research design and methodology, data analysis and writing; NTS, and AI for the research design and funding of seaweed cultivation, and writing. AB also contributed to resource provisioning and software; AS contributed laboratory resources; DC and AG for supervision. All authors have approved the final version of the manuscript.

## Notes

### Competing Interest Statement

The authors have declared no competing interest.

